# A Novel Silver-Ruthenium-Based Antimicrobial Kills Gram-Negative Bacteria Through Oxidative Stress-Induced Macromolecular Damage

**DOI:** 10.1101/2025.01.03.631245

**Authors:** Patrick Ofori Tawiah, Luca Finn Gaessler, Greg M. Anderson, Emmanuel Parkay Oladokun, Jan-Ulrik Dahl

**Author notes:** Address correspondence to Dr. Jan-Ulrik Dahl, School of Biological Sciences, Illinois State University, Campus Box 4120, Normal, IL 61790.

## Abstract

Amplified by the decline in antibiotic discovery, the rise of antibiotic resistance has become a significant global challenge in infectious disease control. Extraintestinal *Escherichia coli* (ExPEC), known to be the most common instigators of urinary tract infections (UTIs), represent such global threat. Novel strategies for more efficient treatments are therefore desperately needed. These include silver nanoparticles, which have been used as antimicrobial surface-coatings on catheters to eliminate biofilm-forming uropathogens and reduce the risk of nosocomial infections. AGXX® is a promising silver coating that presumably kills bacteria through the generation of reactive oxygen species (ROS) but is more potent than silver. However, neither is AGXX®’s mode of action fully understood, nor have its effects on Gram-negative bacteria or bacterial response and defense mechanisms towards AGXX® been studied in detail. Here, we report that the bactericidal effects of AGXX® are primarily based on ROS formation, as supplementation of the media with a ROS scavenger completely abolished AGXX®-induced killing. We further show that AGXX® impairs the integrity of the bacterial cell envelope and causes substantial protein aggregation and DNA damage already at sublethal concentrations. ExPEC strains appear to be more resistant to the proteotoxic effects of AGXX® compared to non-pathogenic *E. coli,* indicating improved defense capabilities of the uropathogen. Global transcriptomic studies of AGXX®-stressed ExPEC revealed a strong oxidative stress response, perturbations in metal homeostasis, as well as the activation of heat shock and DNA damage responses. Finally, we present evidence that ExPEC counter AGXX® damage through the production of the chaperone polyphosphate.

## INTRODUCTION

*Escherichia coli* is characterized by its remarkable diversity: while some members of this species are part of the commensal vertebrate gut microbiota, others are known to cause serious intestinal (i.e. various forms of diarrhea) and extraintestinal diseases (i.e. urinary tract infections [UTIs],bacteremia, pulmonary, skin and soft tissue infections) (1). The most prominent group of extraintestinal *E. coli* (ExPEC) are uropathogenic *E. coli* (UPEC), the etiologic agent of UTIs. UPEC causes 75% of all uncomplicated UTIs (arise spontaneously in otherwise healthy patients) and 65% of all complicated UTIs (refer to various patient-specific factors, such as the presence of a catheter or stent or immunocompromised patients) (2). 405 million UTIs and 267,000 UTI-related deaths worldwide have been estimated in 2019, with women and the elderly population being disproportionally affected (2, 3). For the US, the Centers for Disease Control and Prevention reported 2.9 million emergency department visits and 3.5 million ambulatory visits directly related to UTIs, with associated costs of more than $6 billion (2). UPEC typically reside as commensals in the gut but turn into serious pathogens upon entry into the urinary tract. Planktonically growing UPEC ascend to the bladder, where they must counter various host defense mechanisms, including phagocyte infiltration (4). Further, UPEC are exposed to antimicrobial peptides and oxidants, such as reactive oxygen and chlorine species (ROS/RCS), prior to invading uroepithelial cells to form intracellular biofilm-like communities. Up to 97% of all healthcare-associated UTIs occur in catheterized patients, making catheter-associated UTIs (CAUTIs) a significant problem for hospitalized patients and those living in long-term care facilities (2). Catheterization of patients carries the risk of introducing uropathogens into the bladder lumen, which is often accompanied by a strong immune response and mucosal irritation upon establishment of the infection. Catheters and other medical devices provide an excellent surface layer that pathogens, including UPEC, use for attachment and formation of biofilms, the ultimate cause of CAUTIs (2). The incidence of ExPEC infections in humans has been increasing over the last decade (5), which demonstrates the critical need to better understand the molecular details of their pathogenesis and develop new approaches for prevention and treatment. The need for new treatment regimens becomes even more pressing in light of the rising antibiotic resistance in *E. coli*: symptomatic UTIs are commonly treated with broad-spectrum antibiotics, however, with limited success as indicated by the high number of recurrent UTIs (4, 6, 7). There is also evidence for an increasing number of uropathogens that carry antimicrobial resistance (AMR) genes (2). According to a recent report, UTIs are now the fourth most common cause of deaths related to AMR, with uropathogens representing five out of the six pathogens most commonly associated with AMR-related deaths (8).

Over the last two decades, silver derivates have received increased attention in medical applications, e.g. as antimicrobial surface-coatings on catheters, protecting from biofilm-forming bacteria and reducing the risk of nosocomial infections (9). Silver has long been known for its antibacterial properties, as ancient Greeks used the metal for wound healing (10). Despite its long-standing history and high efficacy against bacteria, the antimicrobial mode of action of silver is poorly understood. Pleiotropic effects have been described and include changes in DNA condensation, membrane alteration, and protein damage. Cationic silver (Ag^+^) interacts with cysteine thiols, destabilizes iron-sulfur clusters, replaces metal-containing cofactors, and elicits ROS production indirectly by a variety of mechanisms, thereby damaging a wide range of proteins (10). Studies with silver ions suggest that Gram-negative bacteria are more susceptible than Gram-positives (11). However, through the rise and spread of AMR, the need for improved surface-coatings on medical devices to eliminate biofilm-forming pathogens via contact-killing has increased (12, 13).

One such promising silver-containing surface-coating is AGXX®, which is comprised of the two transition metals silver (Ag) and Ruthenium (Ru) and is currently only used on waterpipes to prevent bacterial attachment (14, 15). The antimicrobial activity of AGXX® is attributed to its specific coating composition and ability to generate ROS when in contact with organic matter (14, 16). AGXX® was shown to be significantly more potent than classical silver, inhibiting the growth of the Gram-positive bacteria *Staphylococcus aureus* and *Enterococcus faecalis* and eliciting a thiol-specific oxidative stress response (14, 17). Moreover, AGXX® is non-toxic to human cells, and no AGXX® resistance has been reported yet, making it potentially well-suited as a novel antimicrobial surface coating on medical devices (18, 19). However, neither has the antimicrobial efficacy of AGXX® on Gram-negative pathogens been explored nor has the precise mechanism of action of this antimicrobial been investigated. Moreover, how Gram-negative bacterial pathogens protect themselves from AGXX® or mediate the repair of AGXX®-mediated damage has not been studied yet. We recently reported that AGXX® potentiates the cytotoxic activities of aminoglycosides in multidrug-resistant *P. aeruginosa* strains, which is attributed to intracellular ROS accumulation, increased membrane damage, and elevated aminoglycoside uptake (20).

In the current study, we aimed to examine the antimicrobial mode of action of AGXX® alone. We found that the bactericidal effect of AGXX® is indeed primarily based on the formation of various ROS, such as hydrogen peroxide (H_2_O_2_) and superoxide (O_2_^⋅−^), as supplementation of the media with the ROS scavenger thiourea completely abolished AGXX®-induced cell death. We show that AGXX® treatment compromises the integrity of the inner membrane and elicits substantial protein aggregate formation and DNA damage, likely as a result of the ROS production. We also found that AGXX® is more effective on non-pathogenic *E. coli* compared to UPEC strains, indicating that uropathogens have a more efficient defense against AGXX®. Global transcriptomic studies of the AGXX®-stressed UPEC strain CFT073 revealed a strong oxidative stress response and perturbations in metal homeostasis. Furthermore, UPEC exposure to AGXX® resulted in the upregulation of members of the heat-shock and DNA damage response, indicating potential proteotoxic and genotoxic effects of the antimicrobial. Finally, we provide evidence that AGXX®-stressed UPEC generate of the chaperone polyphosphate (polyP) to protect themselves from the proteotoxic effects of AGXX®.

## RESULTS

### AGXX® formulations differ in their antimicrobial potency

Several studies have previously reported about the strong bactericidal effects of the silver-ruthenium-based antimicrobial AGXX® against gram-positive pathogens (14, 16–18, 21, 22). However, whether and to what extent AGXX® compromises Gram-negative bacteria has not been investigated yet. Moreover, the precise mode of action of this antimicrobial has not been elucidated. We recently reported that AGXX® is not only more efficient against the Gram-negative pathogen *P. aeruginosa* than silver, the compound also potentiates the efficacy of aminoglycoside antibiotics (20). Over the recent years, AGXX® has undergone continuous optimization, resulting in a variety of formulations. While these AGXX® formulations all consist of the galvanized silver-ruthenium complex, they differ in various aspects, such as the silver ratio, particle size, and production procedure, with potentially significant consequences for their antimicrobial activity. To compare the effective antimicrobial concentrations of different AGXX® formulations against *E. coli*, we first performed survival analyses and growth inhibition studies of the UPEC strain CFT073 in the presence and absence of AGXX®383, AGXX®394C, AGXX®823, and AGXX®894, respectively. CFT073 cultures in their mid-logarithmic (mid-log) phase were exposed to the indicated concentrations of these AGXX® formulations, and growth (**Supplementary Fig. S1A-D**) and survival (**Supplementary Fig. S1E-H**) were monitored over the defined time intervals. Overall, AGXX®394C (**Supplementary Fig. S1A, E**) was the most potent formulation, which required 5- to 6.3-fold lower concentrations for effective inhibition of UPEC growth and survival compared to AGXX®383, AGXX®894 and AGXX®823, respectively.

### The antimicrobial activity of AGXX® is caused by ROS production

Regardless of the specific AGXX® formulation, the primary antimicrobial action has been proposed to be based on ROS generation although this has not been directly shown yet. To investigate whether AGXX® treatment causes ROS accumulation in Gram-negative bacteria, we utilized generic and ROS-specific redox-sensitive probes for the detection of intracellular ROS in CFT073 cultures grown in the presence and absence of AGXX®. Briefly, we treated exponentially grown CFT073 with the indicated AGXX®394C concentrations for 60 min and quantified intracellular ROS with the 2’,7’-dichlorodihydrofluorescein diacetate (H_2_DCFDA) dye. H_2_DCFDA is a non-fluorescent redox probe that is readily oxidized by different ROS, forming a fluorescent DCF moiety (23). When UPEC was exposed to a sublethal dose (i.e., 30 µg/ml) of AGXX®, we did not detect significant changes in DCF fluorescence, indicating no significant increase in intracellular ROS levels compared to untreated cells **(Fig. 1A)**. However, DCF fluorescence increased 11- to 13-fold upon treatment with 40 µg/ml and 50 µg/ml AGXX® **(Fig. 1A),** two concentrations that were already slightly to moderately bactericidal **(Fig. 1B)**. To examine whether the increase in intracellular ROS contributes to the bactericidal effect of AGXX®, we pretreated CFT073 cultures with a ROS quencher thiourea prior to AGXX® exposure. Thiourea-pretreated UPEC cells showed low DCF fluorescence values despite the subsequent challenge with AGXX®, which were comparable to those observed in untreated cells **(Fig. 1A)**. Likewise, supplementation of thiourea had positive effects on UPEC survival during AGXX® stress, which became even more apparent when the time-killing analysis was performed over 3 hrs **(Fig. 1B, C)**. Overall, our data indicate that AGXX®-induced ROS production is indeed an important part of the compound’s antimicrobial activity. Given the comparatively low sensitivity of H_2_DCFDA and its low specificity towards specific ROS variants, we conducted these experiments with the more sensitive fluorescent probes dihydroethidium (DHE) and Amplex ^TM^ Red, which specifically detect intracellular levels of O_2_^⋅−^ and H_2_O_2_, respectively. Consistent with the data in **Fig 1A**, we observed concentration-dependent increases in O_2_^⋅−^ and H_2_O_2_ of up to 6- and 4-fold after 60 mins of AGXX® exposure starting already at sublethal concentrations (**Fig. 1D & E**). Taken together, our data provide evidence for the direct involvement of various ROS in the bactericidal mode of action of AGXX®, which can be alleviated by the presence of antioxidants.

**Fig 1:**
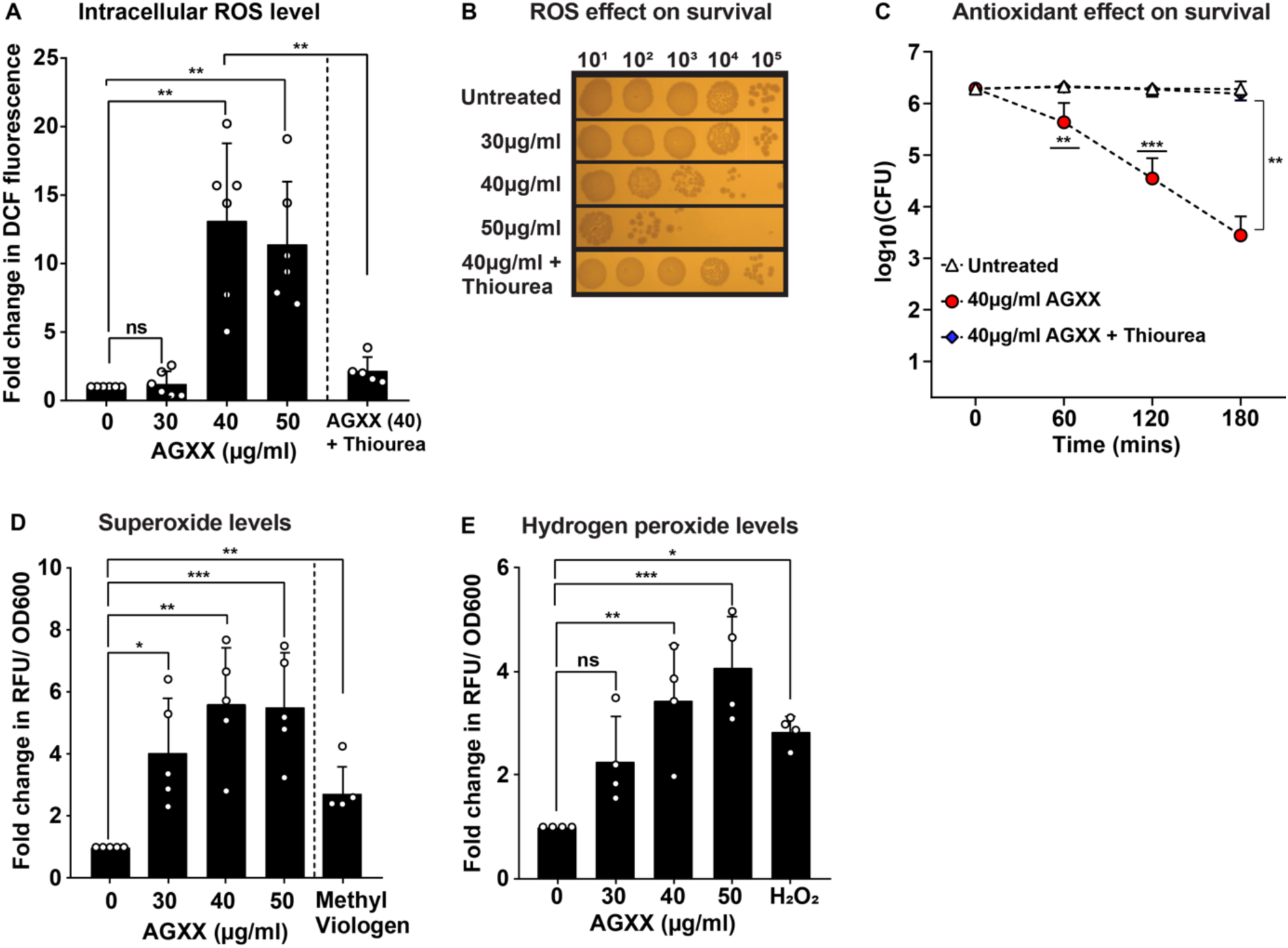
AGXX®-stressed bacteria accumulate large amounts of ROS, which contribute to the bactericidal effects of this antimicrobial. UPEC cells grown to mid-log phase were left untreated or treated with the indicated AGXX®394C concentrations for 60 min before **(A)** intracellular ROS were quantified by H_2_DCFDA, and **(B; C)** survival was determined by serially diluting cells in PBS and spotting 5µl onto LB agar for overnight incubation. **(D)** Intracellular superoxide levels were detected by DHE and **(E)** H_2_O_2_ quantified by Amplex ^TM^ Red. 70 mM thiourea was used to quench ROS; (n= 4-6, ±S.D., one-way ANOVA, Dunnet and Sidak’s multiple comparison test; ns = *P* > 0.05, * *P* < 0.05, ** *P* < 0.01, *** *P* < 0.001).

### AGXX® exposure causes significant membrane damage

The bacterial plasma membrane serves as a permeability barrier that shields the cytoplasmic milieu from harmful compounds. Therefore, damage to the membrane architecture can have severe consequences for bacterial survival. Previous reports of AGXX®-stressed *S. aureus* suggested potential membrane-compromising effects of AGXX®, although this was only based on transcriptional data and lacked direct evidence (16, 17). To investigate whether sublethal AGXX® treatments result in membrane disruptions, we treated exponentially growing CFT073 with the indicated concentrations of AGXX® and examined the integrity of the inner membrane by quantifying propidium iodide (PI) influx into the cell. The inner membrane is not PI permeable if intact due to the fluorophore’s size and charge. However, PI freely enters cells with disrupted membranes, which results in stronger fluorescence values due to the non-specific intercalation of PI with nucleic acids (24). While there was a slight increase in PI uptake in cells challenged with sublethal AGXX® concentrations (i.e. 10-30 mg/ml AGXX®), treatment with 40 µg/ml, a concentration that only caused minimal killing (**Fig. 1B, C),** resulted in ∼150-fold higher PI fluorescence compared to the untreated control (**Fig. 2A**). This correlates well with the fold-change in PI fluorescence observed in UPEC cells exposed to the membrane-targeting antibiotic polymyxin B (PMB) (**Fig. 2A**). However, exposure to sublethal AGXX® concentrations (i.e. 10-30 mg/ml AGXX®) resulted in increased PI fluorescence when challenged for prolonged times (i.e. 120 and 180 min, respectively), indicating a slow acting mechanism of this antimicrobial (data not shown). These findings were further confirmed by live/dead staining. Fluorescence microscopy analyses revealed that while 94% of the untreated cells could only be stained by Syto9, the number of PI-stained cells increased remarkably to 47% when cells had been treated with 40 µg/ml AGXX® (**Fig. 2B; Supplementary Table S1**). Thus, our data provide clear evidence for the membrane-damaging effects of AGXX®, which is likely a result of the AGXX®-induced ROS formation.

**Fig 2:**
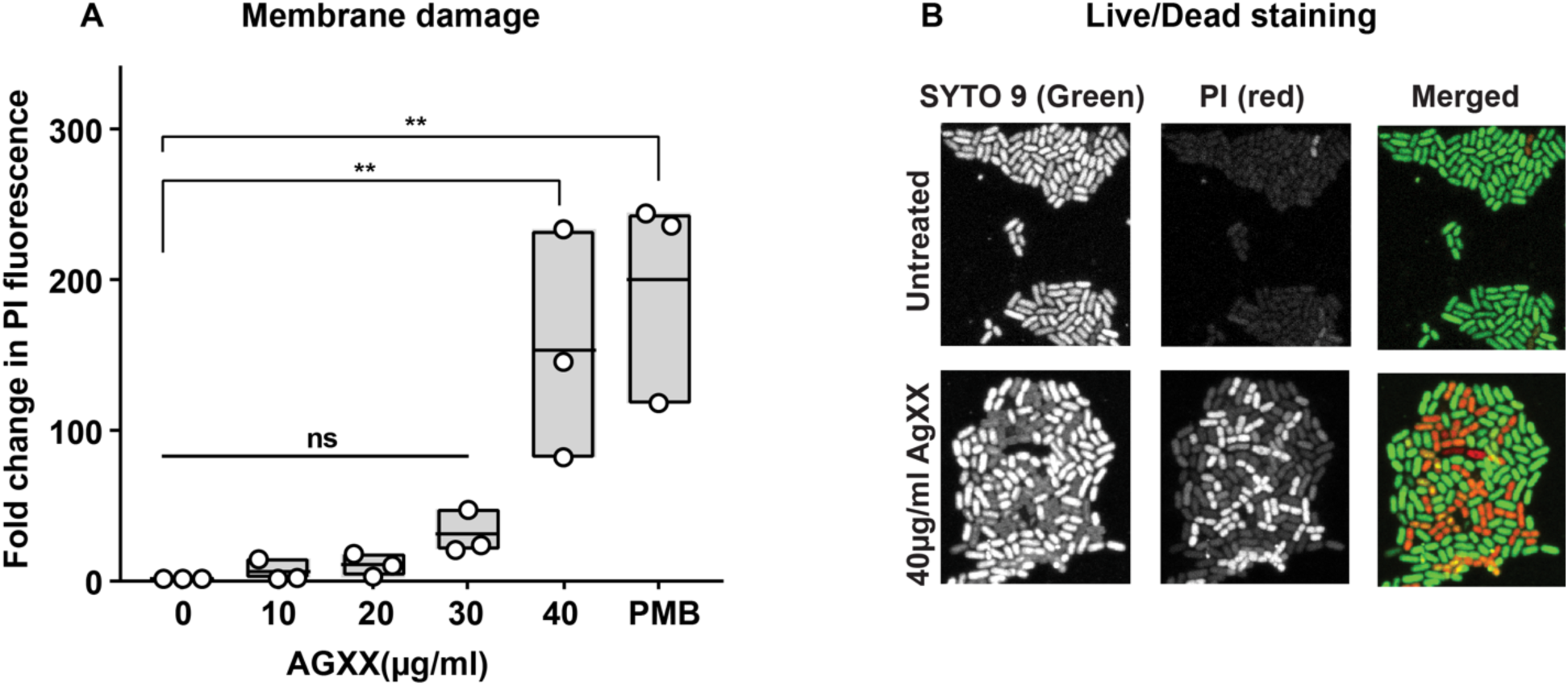
AGXX® stress compromises bacterial membrane integrity. **(A)** CFT073 cells in the mid-log phase were treated with the indicated AGXX®394C concentrations for 60 min, washed in PBS, and stained with 0.5 µM PI. PI fluorescence (λ_Ex/Em_: 535/617nm) was measured by spectrophotometry and normalized to untreated cells (n=3, ±S.D.). **(B)** Samples were washed in PBS after AGXX®394C treatment, incubated with PI/Syto9 in the dark for 15 min at room temperature, and mounted on a glass slide with a 1% agarose pad for 63x imaging using inverted confocal microscopy. One representative image of 4 independent experiments is shown. (one-way ANOVA, Sidak’s multiple comparison test; ns = *P* > 0.05, * *P* < 0.05, ** *P* < 0.01).

### AGXX® causes extensive protein aggregation and increases the cellular demand for molecular chaperones

Previous studies in AGXX®-stressed Gram-positive bacteria reported an elevated transcription of the heat-shock response (16, 17, 22), indicating proteotoxic effects of AGXX®. We decided to monitor AGXX®-induced protein aggregate formation in *E. coli* in real time. We transformed cells with a reporter plasmid that allows for the expression of IbpA fused to sfGFP (IbpA-msfGFP) under the control of the native *ibpA* promoter. IbpA, whose expression is induced under protein unfolding conditions (25, 26), is a universally conserved molecular chaperone that binds unfolded proteins and protein aggregates. Expression of IbpA-sfGFP has been successfully used before to quantify protein aggregation *in vivo* (27). We treated exponentially growing cultures with sublethal AGXX® concentrations for 120 min and determined the cellular sfGFP signal by flow cytometry. AGXX® treatment resulted in a significant intensity shift in sfGFP fluorescence, suggesting that AGXX®-mediated damage triggers IbpA expression (**Fig. 3A**). Pretreatment with the ROS quencher thiourea eliminated any AGXX®-induced sfGFP fluorescence back to the levels of untreated cells (**Fig. 3A**). Cell samples that were collected before and after treatment with sublethal AGXX® concentrations for 90 min were also embedded onto agarose pads on a glass slide for visualization and quantification of the green IbpA-msfGFP foci under the fluorescence microscope. Given that protein aggregation occurs naturally even in non-stressed cells, it was not surprising to see 30% of the untreated cells with one IbpA-sfGFP foci per cell (**Fig. 3B, C**). In contrast, treatment with sublethal (i.e., 32 and 37.5 µg/ml) and slightly bactericidal AGXX® concentrations (i.e., 40 µg/ml; **Fig 1B, C**) resulted in a significant, concentration-dependent increase in IbpA-sfGFP foci formation. In fact, ∼80% of the cells showed IbpA-sfGFP foci formation, with the majority of cells containing more than one foci per cell (**Fig. 3B, C; Supplementary Table S2**). Overall, these data show that AGXX® stress impairs protein homeostasis, which results in the induction of the expression of heat shock proteins to cope with the negative consequences of AGXX® stress.

**Fig 3:**
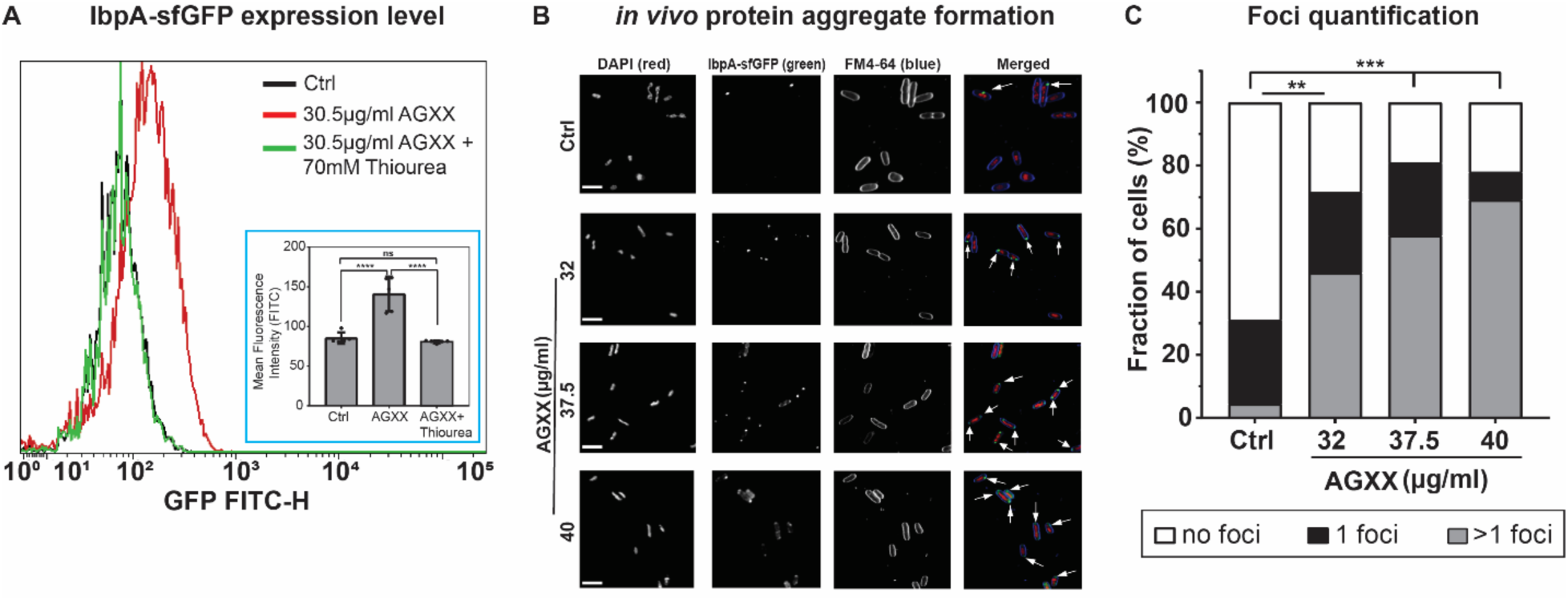
AGXX® causes extensive protein aggregation and increases the cellular demand for molecular chaperones. **(A)** Cellular IbpA-sfGFP fluorescence was monitored via flow cytometry after exposure of *E. coli* to sublethal AGXX®394C treatment for 120 min; (n=5, ±S.D.). One representative image of five independent experiments is shown. **(B)** Exponentially growing cells were either left untreated or treated with sublethal AGXX®394C concentrations for 90 min. Samples were harvested, washed with PBS, and incubated with DAPI (nucleic acid stain) and FM4-64 (membrane stain) in the dark for 15 min. Cells were mounted on a glass slide with a 1% agarose pad for imaging at 63x via inverted confocal microscopy. Arrows illustrate foci formed when IbpA binds to protein aggregates in *vivo*. One representative image of 4 independent experiments; [scale bar: 7.5 µm]. **(C)** Confocal images were quantified by counting IbpA-sfGFP foci (n=4) (one-way ANOVA, Sidak’s multiple comparison test; two-way ANOVA, Tukey’s multiple comparison test (compare total foci in untreated to AGXX treatment); ns = P > 0.05, * *P* < 0.05, ** *P* < 0.01, *** *P* < 0.001, **** *P* < 0.0001).

### AGXX® is genotoxic, resulting in DNA double-strand breaks

We have previously reported about the synergistic effects between AGXX® and aminoglycoside antibiotics, rendering drug-resistant *P. aeruginosa* strains sensitive again (20). The bactericidal effect of this synergy is mediated by a significant increase in outer and inner membrane permeability, which exacerbates aminoglycoside influx into the cell. Intriguingly, transcriptomic data revealed an upregulation of the DNA damage response in *P. aeruginosa* exposed to a combination of AGXX® and aminoglycosides (20). We suspect this effect to be caused by AGXX® as aminoglycosides specifically target protein translation and have no known detrimental effects on DNA (20). To examine a potential genotoxic activity of AGXX®, we first monitored the transcript levels of *sulA,* a hallmark gene of the bacterial DNA damage response, in UPEC cells that were treated only with AGXX®. Intriguingly, s*ulA* mRNA levels were ∼8-fold upregulated upon treatment with sublethal AGXX® concentrations for 30 min (**Fig. 4A**). We then sought to directly determine the extent to which AGXX® treatment causes DNA damage. We recombinantly expressed the bacteriophage protein Gam, which specifically binds to DNA double-strand breaks in mammalian and bacterial cells (28), fused to the monomeric superfolder green fluorescent protein (Gam-msfGFP) in *E. coli* to detect DNA double-strand breaks. We treated exponentially growing cells for 3 hrs with the indicated concentrations of AGXX®, prepared samples for visualization by fluorescence microscopy, and quantified Gam-sfGFP bound to DNA double-strand breaks, which appear as green, fluorescent foci. Under non-stress conditions, only ∼10% of all cells contained detectable Gam-sfGFP foci (**Fig. 4B, C; Supplementary Table S3**). In contrast, treatment with sublethal (i.e., 37.5 mg/ml) and slightly bactericidal concentrations (i.e., 40 mg/ml) significantly increased the number of cells with DNA double-strand breaks to over 40%, which was comparable to treatments with sublethal concentrations of the DNA damaging fluoroquinolone ciprofloxacin (Cpx) (**Fig. 4B, C; Supplementary Table S3**). Cells with more than one foci increased ∼6-fold upon AGXX® exposure, highlighting the severity of DNA damage induced by AGXX®. Additionally, we observed that some of the AGXX-stressed cells appear filamentous, which has been reported as a typical DNA damage response, particularly as a result of the activation of the SOS response (29).

**Fig 4:**
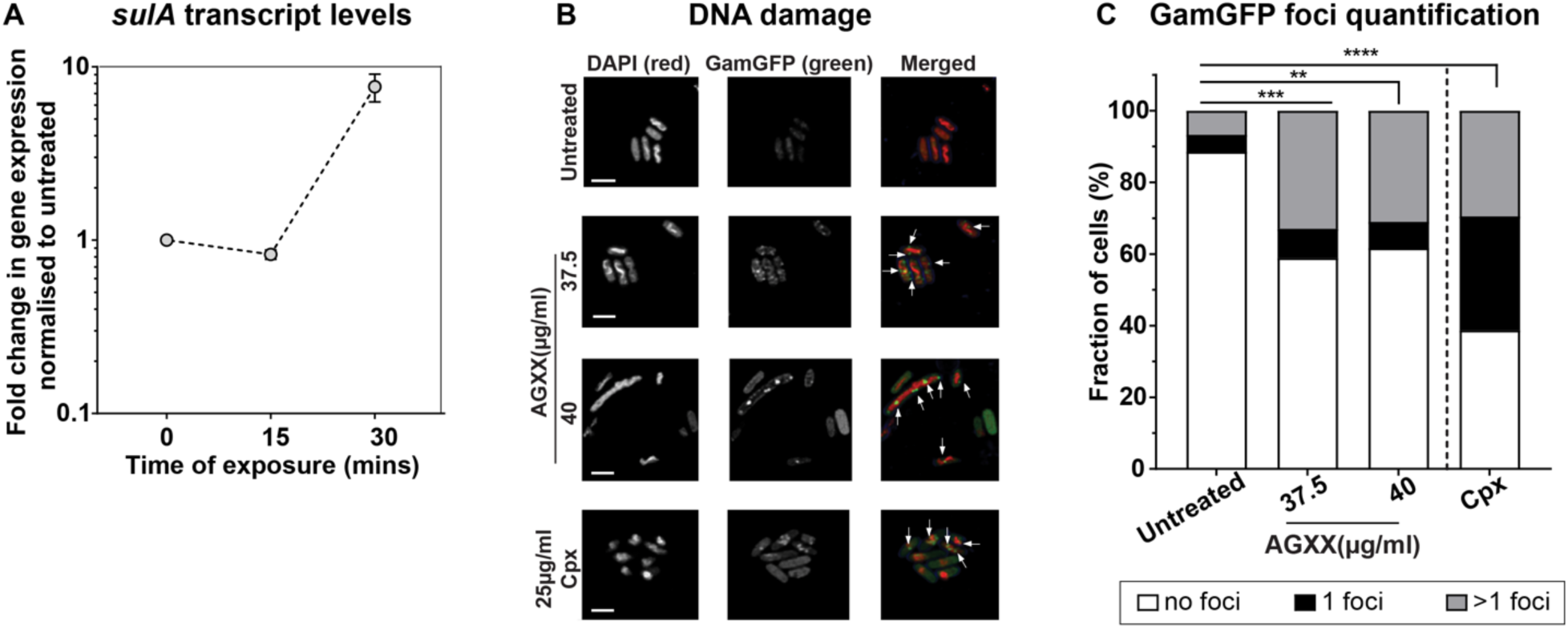
AGXX® causes DNA double-strand breaks. **(A)** *sulA* mRNA levels of AGXX®394C-treated UPEC CFT073 cells were determined by qRT-PCR. Transcript levels were normalized to the housekeeping gene *rrsD* and calculated as fold-changes based on the expression levels in the untreated control (n = 3, ±S.D.). **(B)** Cells expressing Gam-sfGFP were exposed to AGXX®394C for 3 hours, washed, incubated with DAPI in the dark for 15 min, and visualized by confocal microscopy. Ciprofloxacin was used as a positive control. Arrows indicate Gam-sfGFP foci on the DAPI-stained DNA. One representative image of three independent experiments is shown [Scale bar: 5 µm]. **(C)** Gam-sfGFP foci were quantified by counting the number of foci per cell (n=3); [two-way ANOVA, Tukey’s multiple comparison test (compare total foci in untreated to AGXX® treatment); ns = *P* > 0.05, * *P* < 0.05, ** *P* < 0.01, *** *P* < 0.001].

### *E. coli* pathotypes differ in their resistance to AGXX®

We recently reported that *E. coli* pathotypes differ in their resistance towards RCS; while enteropathogenic and non-pathogenic K-12 strains were equally sensitive, members of the UPEC pathotype tolerated significantly higher levels of these neutrophilic oxidants (30). UPECs increased RCS resistance is the result of a specific inactivation of an RCS-sensing transcriptional repressor that causes increased expression of *rcrB* (30). The gene is part of one of UPECs pathogenicity islands, does not exist in *E. coli* K-12 strains, and encodes the putative membrane protein RcrB, the precise function of which is still under investigation. Comparative genomics revealed that more than 80% of open reading frames in three of the most prominent UPEC lab strains are identical, yet only 37% of their genomes also exist in commensal *E. coli* (31). Many of the UPEC-specific virulence factors have been linked to their ability to persist in the urinary tract despite aggressive host defense mechanisms (32–34). To test whether UPEC also tolerates AGXX® more efficiently, we compared the impact of AGXX® on the growth and survival of non-pathogenic K-12 strain MG1655 and the UPEC strain CFT073. Both strains were grown to mid-log phase and either left untreated or treated with increasing concentrations of AGXX®. The presence of AGXX® completely stopped growth and significantly impaired survival of MG1655 at concentrations that had no effect on CFT073 (**Fig. 5A, B**). Thus, out data indicate that UPEC has evolved specific strategies to better deal with the negative consequences of AGXX® stress, which non-pathogenic *E. coli* lack. Next, we sought to determine whether UPECs superior tolerance towards RCS and AGXX® is mediated by a common denominator, namely the expression of the putative membrane protein RcrB (30, 35). Quantitative reverse transcriptase PCR (qRT-PCR) revealed that hypochlorous acid (HOCl), the most potent RCS, induces transcription of *rcrB* whereas no significant changes in *rcrB* mRNA level were detected upon treatment with AGXX® (**Fig. 5C**). Likewise, we did not observe significant RcrB-dependent differences in survival when UPEC cells with and without RcrB were exposed to AGXX® (**Fig. 5D**), excluding the possibility that RcrB is responsible for UPECs superior AGXX® resistance.

**Fig. 5:**
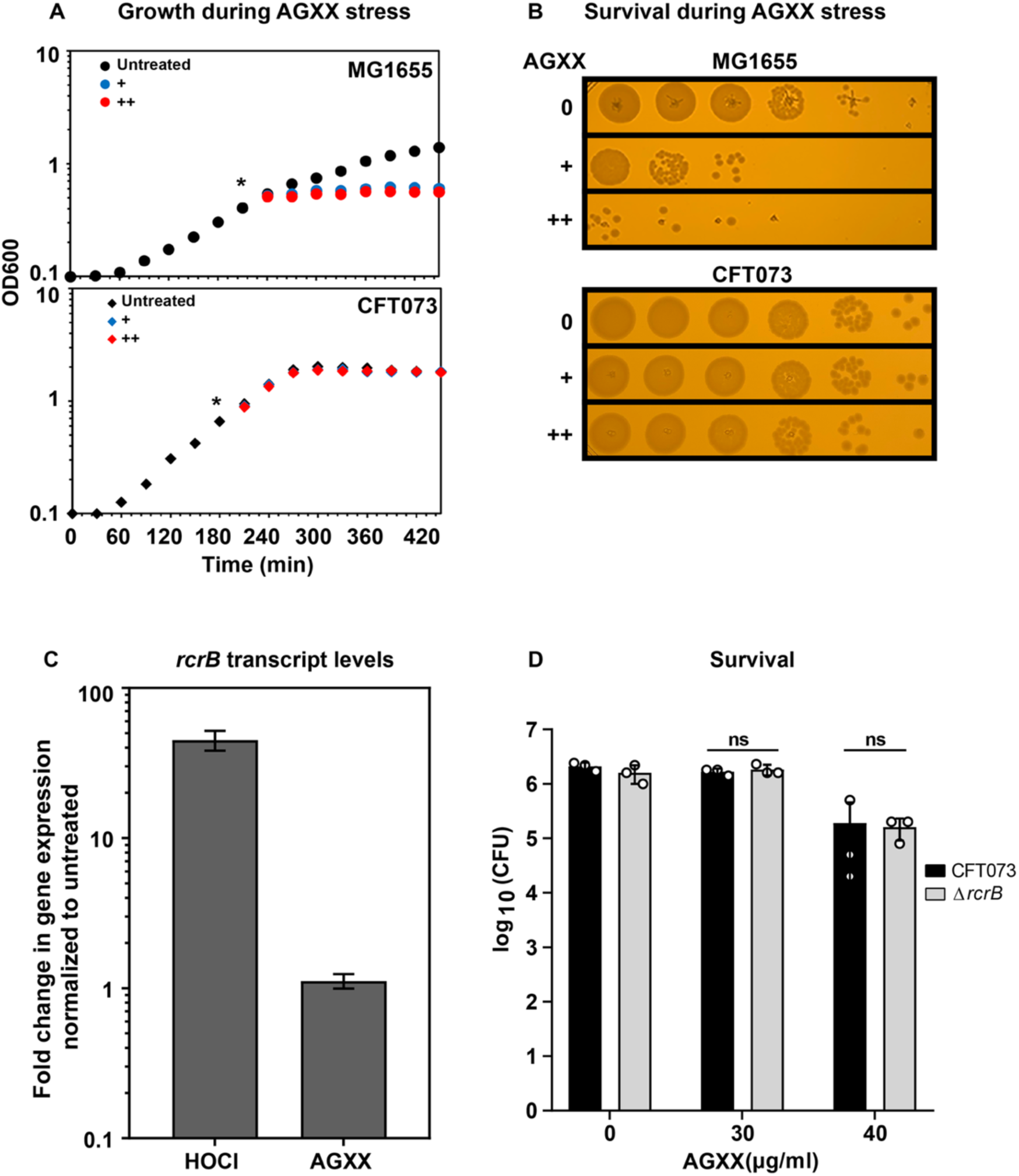
UPEC strain CFT073 shows increased tolerance to AGXX® compared to K-12 *E. coli* strain MG1655, which is independent of RcrB. Overnight cultures of *E. coli* strains MG1655 and CFT073 were diluted into MOPSg to an OD_600_∼0.1 and grown to mid-log phase (OD_600_∼0.5). Cells were either left untreated or exposed to increasing concentrations of AGXX®. Growth **(A)** and survival **(B)** were recorded. **(A)** AGXX® concentrations that resulted in a growth arrest of MG1655 had no effect on CFT073. **(B)** After 60 min of AGXX exposure, cells were serially diluted and spot-titered onto LB agar plates for overnight incubation at 37 °C. One representative image of at least three independent experiments with similar outcomes. **(C)** *rcrB* mRNA levels of HOCl and AGXX®-treated UPEC CFT073 cells were determined by qRT-PCR after 15 mins. Transcript levels were normalized to the housekeeping gene *rrsD* and calculated as fold changes normalized to that of untreated control (n = 3, ±S.D.). **(D)** Exponentially grown CFT073 WT and Δ*rcrB* cultures were either left untreated or treated with AGXX®. After 60min, cells were serially diluted and spot-titered onto LB agar plates for overnight incubation at 37 °C. (n = 3, ±S.D.).

### AGXX®-induced protein aggregation is less pronounced in members of the UPEC pathotype

Our data show that AGXX® induces extensive protein unfolding and aggregation (**Fig. 3**), likely due to its ability to readily oxidize redox-sensitive amino acid side chains, which shifts the equilibrium of proteins more toward their aggregation-prone state (17). AGXX®-induced protein aggregation explains the upregulation of members of the heat shock regulon, such as *ibpA* (**Fig. 3A**), whose expression is triggered by the accumulation of unfolded proteins (36). To determine whether UPECs superior tolerance to AGXX® is due to its improved ability to deal with AGXX®-mediated protein aggregation, we compared changes in transcript levels of the three molecular chaperone genes *ibpA, ibpB*, and *dnaK* between AGXX®-sensitive K-12 strain MG1655 and UPEC strain CFT073. All three genes were significantly upregulated in MG1655 at AGXX® concentrations that had little to no effect on their transcript levels in CFT073 (**Fig. 6A**). Notably, these concentrations significantly inhibited MG1655 growth and survival but did not compromise CFT073 (**Fig. 5**). At higher AGXX® concentrations, mRNA levels were also induced in CFT073 (**Fig. 6A**), indicating that the pathogen indeed depends on a functional heat-shock response but does so only at comparatively higher AGXX® concentrations. Next, we sought to directly compare AGXX®-mediated protein aggregation in both strains following our recently published assay (37). Cells were either left untreated or treated with the indicated AGXX® concentrations for 45 min, lysed, and protein aggregates separated from soluble proteins. The isolated protein fractions were then separated by SDS-PAGE and visualized by Coomassie staining. We observed a substantially greater extent of protein aggregates in AGXX®-treated MG1655, which correlates well with the inability of this strain to cope with these AGXX® concentrations, all while CFT073 remained unaffected (**Fig. 6B**). Visualization of the soluble protein fraction revealed an AGXX®-induced decrease for both strains. Overall, our data indicate that the proteotoxic effect of AGXX occurs in both strains but is better tolerated by UPEC.

**Fig 6:**
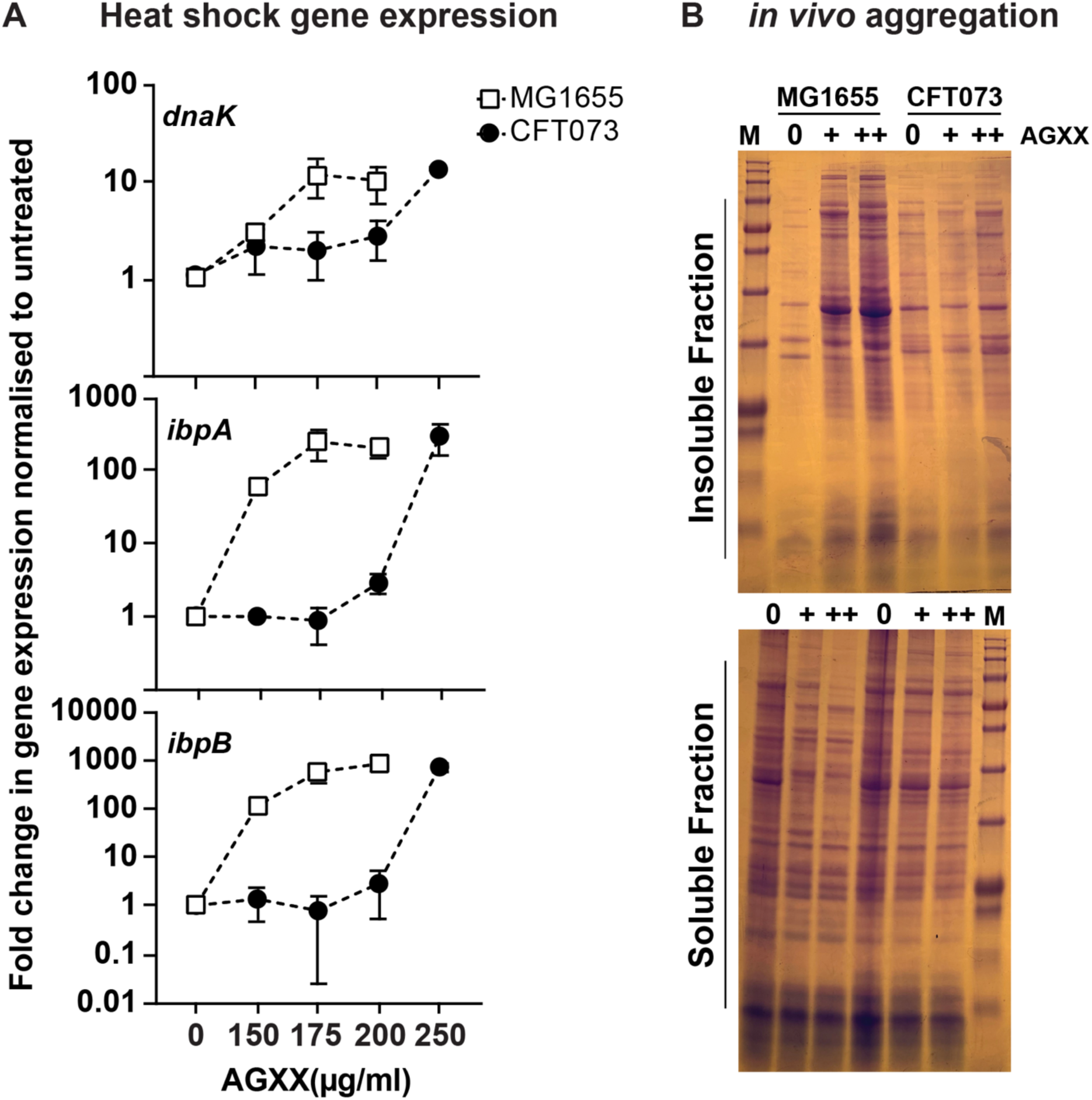
AGXX®-induced protein aggregation is less pronounced in members of the UPEC pathotype. Mid-log phase cultures of K-12 strain MG1655 and UPEC CFT073 were exposed to the indicated AGXX®823 concentrations for 30 min. **(A)** Total RNA was extracted, genomic DNA removed, and mRNA reverse-transcribed into cDNA. qRT-PCR analysis was performed for differential expression analyses of genes *ibpA, dnaK*, and *ibpB*, which were normalized to the housekeeping gene *rrsD* and the untreated cells; (n=5-7, ±S.D.) **(B)** The extent of protein aggregation was determined after harvesting and cell lysis. Protein aggregates and soluble proteins were separated, extracted, separated by SDS-PAGE, and visualized by Coomassie staining. One representative image of 5 biological replicates is shown.

### AGXX® elicits widespread transcriptional changes in UPEC

Bacteria have acquired a multitude of mechanisms to respond rapidly to changes in their environment, including the tight control of gene expression. Our UPEC survival studies (**Supplementary Fig. S1**) suggest a more slow-acting killing mechanism for AGXX® compared to fast-acting oxidants such as HOCl, which causes changes in gene expression very rapidly and kills bacteria in as little as a few minutes (35, 38, 39). To monitor changes in mRNA level during AGXX® over time, we performed a time-course experiment focusing on the three select heat-shock genes *ibpA, ibpB*, and *dnaK*, as previous studies in Gram-positive bacteria suggested that AGXX® induces members of the heat-shock response (16, 22). We challenged mid-log phase CFT073 cells with sublethal concentrations of AGXX® and determined the transcript levels of the three genes at the indicated time points. We did not detect any substantial changes in mRNA levels of any of the genes after 15 min of treatment (**Supplementary Fig. S2**), indicating that AGXX® indeed elicits a slower response. However, after 30 min of AGXX® treatment, mRNA levels of all three genes increased 10- to 100-fold and remained highly upregulated over the next 30 min (**Supplementary Fig. S2**).

Next, we conducted RNAseq analysis to globally monitor changes in CFT073 gene expression in response to sublethal AGXX® stress. CFT073’s chromosome shows a mosaic structure in the distribution of backbone genes, which are shared with MG1655, and “foreign” genes that presumably have been acquired horizontally (33). We, therefore, reasoned that any CFT073 genes that are highly induced upon AGXX® stress but absent in K-12 strains, such as MG1655, could contribute to CFT073’s elevated AGXX® resistance. For the transcriptome analysis, we compared the expression values of the stress-treated cells to non-stress-treated controls. We set a false discovery rate (FDR) of <0.05 as a threshold for significance and considered transcripts as upregulated when they showed a log_2_ fold change of >1.5 and downregulated when they showed a log_2_ fold change of <-1.5. AGXX® treatment of CFT073 resulted in the upregulation of 179 genes, 121 of which were ≥ four-fold induced. 40% of the upregulated genes (3 to 49-fold induced) are uncharacterized and have no biological function associated, of which 25% are UPEC-specific (**Fig 7A; Supplementary Table S4&S5**). 63 genes were downregulated, of which 31 transcripts were at least four-fold reduced under AGXX® stress. 24 of the downregulated genes were also uncharacterized, of which 9 were UPEC-specific (**Fig 7A; Supplementary Table S4&S5**). Most notably, genes associated with the bacterial oxidative stress response were highly upregulated, including *soxS, sodA, nemA, ahpF, trxC, ybbN,* and *grxA* (**Fig. 7A**; see purple IDs), confirming that AGXX®-treated UPEC experience significant oxidative stress. Furthermore, several genes encoding proteases and molecular chaperones were significantly elevated in AGXX®-stressed UPEC (**Fig. 7A**; see green IDs), confirming the proteotoxic effects of this antimicrobial. AGXX® treatment appears to also affect metal ion homeostasis given the highly induced expression of numerous copper-responsive (see blue IDs) and iron-sulfur cluster biosynthesis genes (e.g., *iscXA*) (**Fig. 7A; Supplementary Table S4**). Notably, mRNA levels of *mutM,* which encodes DNA glycosylase, were also highly upregulated, indicating a response to oxidative DNA damage (40). Among the significantly repressed genes in AGXX®-stressed CFT073 were genes encoding proteins involved in curli assembly (i.e., *csgEDF*), outer membrane proteins (i.e., *ompF, nmpC* and *c2348*), respiratory transport chain complex I (*nuoABCEGH)* and iron uptake systems (i.e., *ycdO, ycdB*), respectively. For a more detailed understanding of the AGXX®-induced transcriptional changes, we grouped the differentially expressed gene according to their corresponding gene ontology term for biological processes using the KEGG database (41). The majority of the differentially expressed genes in AGXX®-stressed CFT073 cells are associated with signaling and cellular processes (e.g., signal transduction, defense mechanisms, secretion), metabolism (e.g., lipid, carbohydrates, amino acids biosynthesis), transcription and translation (i.e., transcriptional regulators, mRNA, tRNA biogenesis) and protein quality control (e.g., chaperones, proteases) (**Fig 7B)**. A large number of the differentially expressed genes were of unknown function. In summary, our data indicate that AGXX® causes a strong oxidative stress response, disrupts metal homeostasis, and induces the expression of genes involved in protein and DNA damage repair.

**Fig 7:**
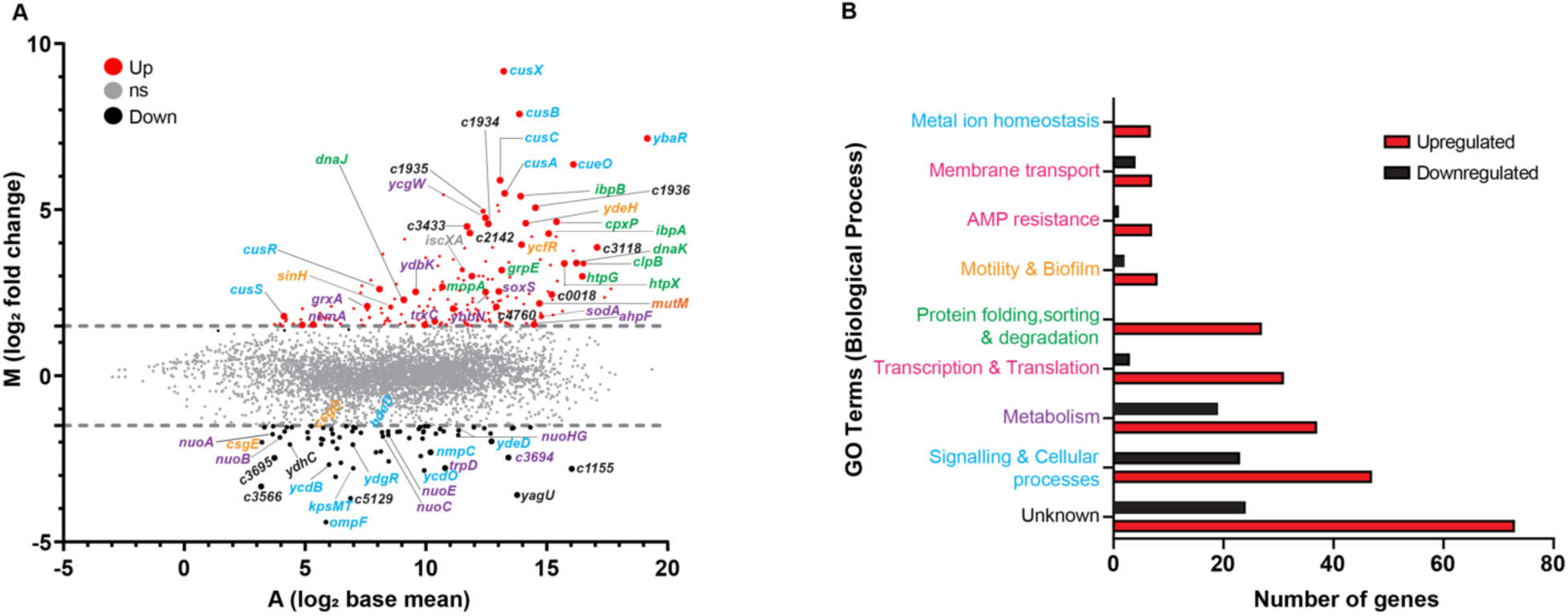
AGXX® exposure of UPEC elicits significant changes in global gene expression. **(A)** Exponentially growing CFT073 cells were incubated with a sublethal concentration of AGXX®394C for 30 min. Transcription was stopped by the addition of ice-cold methanol. Reads were aligned to the CFT073 reference genome (accession number: AE014075). Data are visualized as a ratio/intensity scatter plot (M/A-plot) of differentially expressed genes in AGXX®-treated CFT073 cells. Statistically significantly upregulated genes are depicted above the blue dashed line, whereas statistically significantly downregulated genes are presented as black dots below the black dashed line (*M* ≥1.5 or ≤ -1.5, *P*≤ 0.05). Light gray dots represent genes with no significant fold change in transcript level upon AGXX® treatment (*P*> 0.5). Many of the upregulated genes can be categorized into metal ion homeostasis (blue IDs), protein homeostasis (green IDs), DNA damage (orange ID), and oxidative stress response (purple IDs), respectively. Transcriptome analysis was performed from three independent biological replicates. **(B)** Number of significantly differentially expressed genes of AGXX-stressed CFT073 grouped based on GO terms for biological processes. Genes with more than one biological process were assigned to their respective GO term from KEGG pathway database.

### Polyphosphate protects UPEC from AGXX® stress

Previous studies by us and others have identified that UPEC strains with compromised polyphosphate (polyP) synthesis are more sensitive to RCS (42) and elevated temperatures (43), as well as show impaired biofilm (42, 44) and persister cell formation (42, 45). Although the molecular function of polyP remains enigmatic for most of these phenotypes, polyP has been identified as a chemical chaperone that heat- or RCS-stressed bacteria produce to protect their proteome from aggregation (43, 46, 47). While polyP is highly conserved and has been detected in all three domains of life, only in bacteria have the enzymes of polyP metabolism been well characterized. The generation of polyP_(n+1)_ is reversibly catalyzed by polyP kinase (Ppk) enzymes that reversibly transfer a terminal phosphate of ATP to a growing chain of polyP_(n)_ (48, 49). To assess whether polyP protects UPEC from AGXX® stress, we compared the impact of AGXX® treatments on the growth and survival of UPEC cells with and without functional polyP synthesis (i.e., WT vs. Δ*ppk*=ΔpolyP). Strains were grown in MOPSg media and treated with the indicated AGXX® concentrations in the early exponential phase (OD_600_= 0.3-0.35), stationary phase (OD600∼ >2.0), and stationary phase cells that were diluted back to OD_600_=0.35, respectively. Samples for survival analyses were taken after 180, 240, and 150 min, respectively. Independent of the growth phase, UPEC cells lacking the ability to produce polyP showed a two- to four-log reduction in survival compared to WT cells, suggesting that polyP production is highly beneficial to UPEC during AGXX® stress (**Fig. 8A-C**). Given that polyP-deficient cells also showed increased susceptibility towards silver nitrate and H_2_O_2_ (**Supplementary Fig. S3**), it remains unclear whether the metal or ROS stimulates the production of this bacterial stress defense system. By examining the area under the growth curve, which inversely correlates with an increased lag phase and indicates enhanced growth inhibition, we observed a similar trend between both strains in growth curve-based assays: while WT cells were able to grow in the presence of 2.9- and 3.1 µg/ml AGXX®, these concentrations were highly inhibitory to the ΔpolyP strain. Interestingly, the addition of exogenous polyP completely rescued the increased AGXX® sensitivity of polyP-deficient cells, restoring their growth to WT level (**Fig. 8D**). In summary, these data support a protective role of polyP under AGXX® stress.

**Fig 8:**
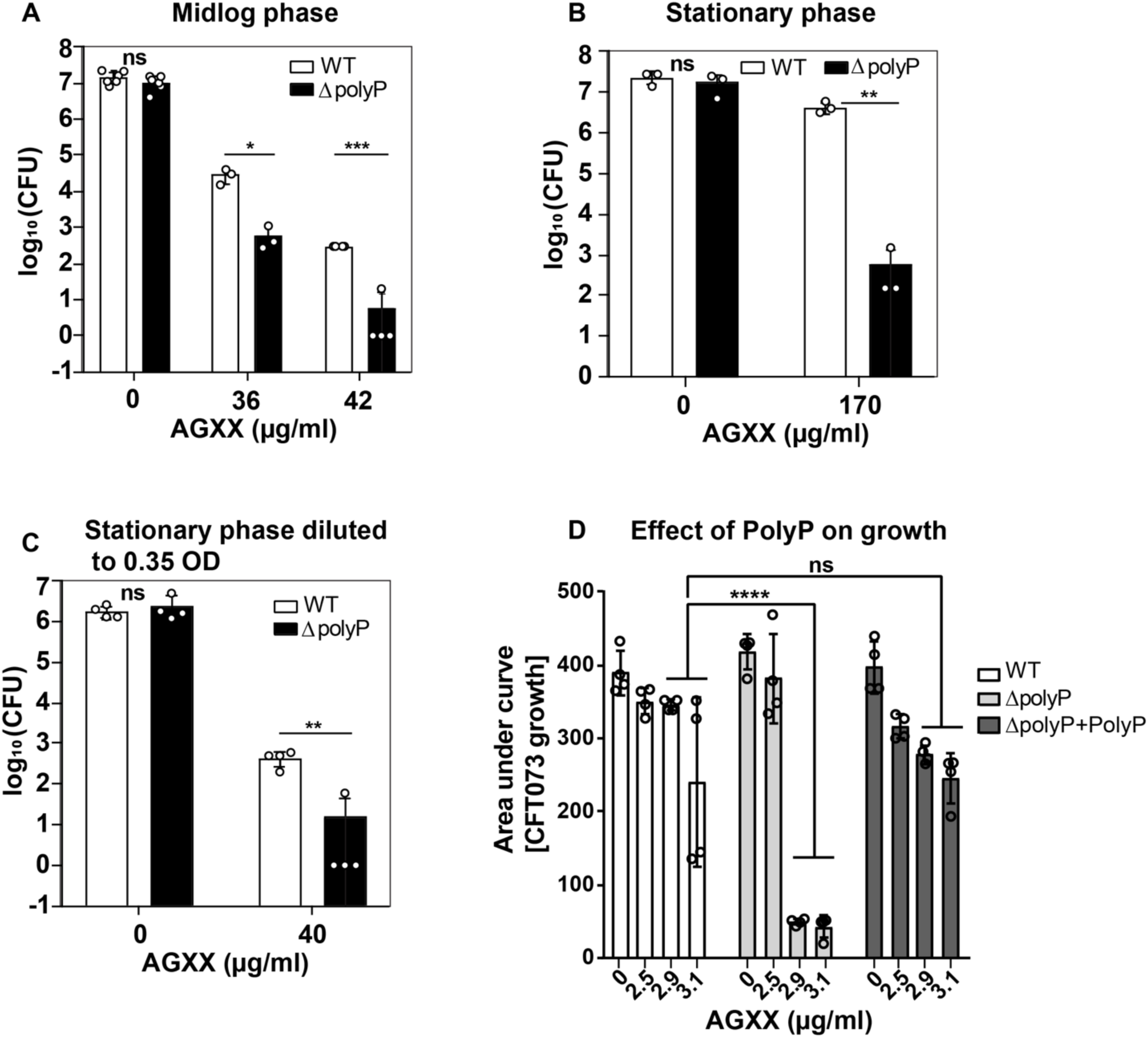
Polyphosphate protects UPEC from AGXX® stress. The role of polyP for UPEC growth and survival during AGXX®394C stress was determined in cells of the **(A)** exponential phase, **(B)** stationary phase, and **(C)** stationary phase cells diluted back into fresh MOPsg to OD_600_=0.35. After 180, 240, and 150 min, samples were serially diluted in PBS, spot-titered on LB agar, and incubated for 20 hrs for CFU counts (n= 3-6, ±S.D.). (D) WT, ΔpolyP, and ΔpolyP supplemented with 4 mM PolyP cultures were cultivated in MOPSg media in the presence of the indicated AGXX® concentrations. Growth was monitored at 600 nm for 16h and calculated as the area under the growth curve; (n=4, ±SD; student t-test; ns = *P* > 0.05, * *P* < 0.05, ** *P* < 0.01, *** *P* < 0.001, **** *P* < 0.0001.)

## DISCUSSION

Previous studies of the novel silver-containing coating AGXX® revealed its strong bactericidal and anti-biofilm effects against Gram-positive bacteria, such as *E. faecalis* and *S. aureus* (14, 16–18, 21, 22). We now provide evidence that AGXX® is also effective against Gram-negative bacteria, such as ExPEC. The bactericidal effects of AGXX® are primarily based on its ROS-producing capabilities as the presence of antioxidants completely abolished AGXX®-induced cell death. While previous reports have already shown the proteotoxic effects of AGXX® in studies with Gram-positive pathogens, we demonstrated for the first time AGXX®-induced protein aggregate formation in living bacterial cells. Moreover, our studies are the first to show that AGXX® treatment compromises the integrity of the inner membrane and elicits substantial DNA damage. Compared to the non-pathogenic *E. coli* K-12 strain MG1655, ExPEC strain CFT073 showed improved tolerance towards the proteotoxic effects of AGXX®. Our global transcriptomic studies of AGXX®-stressed CFT073 revealed a strong oxidative stress response and perturbations in metal homeostasis. Additional signatures of the ExPEC’s transcriptional response to AGXX® stress include the induction of the heat shock and DNA damage responses, which aligns well with the increased protein aggregation and DNA damage observed in AGXX®-treated *E. coli*. ExPEC responds to AGXX® stress by producing the chemical chaperone polyphosphate (polyP) as a defense strategy, likely to prevent AGXX®-induced protein aggregation.

### AGXX® formulations differ in their antimicrobial activities

A comparison of the effects of four different AGXX® formulations (i.e. 383, 394C, 823, and 894) on the growth and survival of ExPEC revealed differences in their antimicrobial efficacies. AGXX® formulations characterized by a smaller particle size showed increased bactericidal activity and required lower concentrations for effective bacterial killing. This was particularly noticeable for the AGXX® formulations 394C and 823 with particle sizes between 1.5 and 2.5 µM, which provide a larger surface area-to-volume ratio than formulations 894 and 383. A larger surface area-to-volume ratio, in turn, increases the likelihood of contact killing due to sufficient contact between the bacterial cell and the AGXX® coating (50). Similar correlations between particle size and antimicrobial activity have been made in previous studies using metal nanoparticles (NP) such as AgNPs (51), zinc oxide (52), and 4,6- diamino-2-pyrimidine thiol-capped gold nanoparticles (53). Likewise, comparisons of the antimicrobial potency of different AGXX® formulations against *S. aureus* revealed more efficient killing by AGXX®373 compared to 383, a formulation that is characterized by a smaller particle size (18).

### AGXX®’s primary antimicrobial activity relies on ROS production

While the antimicrobial effects of silver and silver-containing agents have been studied for a long time, their mechanism of action remains poorly understood (9, 54–58). The noble metal has been proposed to cause pleiotropic effects, including protein mis-metalation, DNA damage, and imbalanced redox homeostasis, which can overburden protective bacterial defense mechanisms and result in bacterial cell death (10, 59, 60). The antimicrobial effects of AGXX®, on the other hand, have been proposed to rely exclusively on ROS formation, which is based on several transcriptome studies (16, 17, 21, 22). Moreover, using spectroscopic methods, Clauss-Lendzian *et al*. provided direct evidence for the formation of H_2_O_2_, when *E. faecalis* was exposed to AGXX®. However, whether the amounts of H_2_O_2_ produced were sufficient to kill the pathogen was not studied (16). Using specific ROS-sensitive fluorophores, we confirmed that AGXX® produces significant amounts of ROS when in contact with cells (**Fig. 1**). Besides a significant increase in H_2_O_2_ levels, we have also detected an AGXX®-mediated increase in superoxide production. We propose that superoxide is likely the main product of AGXX®-mediated ROS production, which then may further be reduced to H_2_O_2_. Notably, ExPEC cells even accumulated high levels of ROS upon exposure to only sublethal AGXX® concentrations indicating a severely imbalanced redox homeostasis during AGXX®-stress. This conclusion is also supported by our RNAseq analysis, which showed several genes encoding antioxidant systems among the highly induced hits in AGXX®-stressed ExPEC (**Fig. 7**). However, while oxidatively stressed bacteria can manage ROS accumulation to some extent, exceeding a certain threshold of stressor concentration overburdens the cellular stress response machinery and results in cell death due to the severe consequences of oxidative damage (61, 62). Our study is also the first to provide direct evidence that ROS production is the main mode of killing for AGXX®, given that the use of the ROS quencher thiourea completely abolished its antimicrobial effects. This is in alignment with our previously reported observation that the synergy between AGXX® and aminoglycoside antibiotics relies on the availability of molecular oxygen, which was almost completely abolished when cells were grown under anaerobic conditions (20). Overall, our data point to a major role of ROS in the killing mode of AGXX® in bacteria. The transition metals silver and ruthenium were proposed to form an electric field leading to a reduction of molecular oxygen and subsequent ROS formation (14). AGXX®’s oxidative stress mode of action is further supported by transcriptome analyses of several Gram-positive bacteria since many genes encoding important antioxidant systems such as thioredoxins, catalase, superoxide dismutase, and glutathione synthetase were highly upregulated (16, 17, 21, 22). Redox biosensor measurements in *S. aureus* strain USA300 as well as studies of the low molecular weight thiol bacillithiol further support a thiol-reactive mode of action of AGXX® (17).

### AGXX® treatment causes significant membrane damage, protein aggregation, and DNA damage

Several independent studies in Gram-positive bacteria, which were primarily based on transcriptomic approaches, have pointed towards potential proteotoxic effects of AGXX® (16, 17, 21, 22). The most direct evidence, presented by two independent studies, revealed significant protein aggregation in bacterial cell lysates that were treated with AGXX® (17, 37). To provide more direct evidence of cytoplasmic proteins indeed being the primary targets of AGXX®, we endogenously expressed the fluorescently labelled biosensor IbpA from the chromosome of *E. coli* to visualize and quantify AGXX®-mediated protein aggregation in living cells. As part of the bacterial heat shock response, molecular chaperones such as IbpA protect the vulnerable proteome from irreversible aggregation (63). These important stress response proteins bind unfolded and misfolded proteins to prevent their aggregation until non-stress conditions are restored that allow for protein refolding by ATP-dependent chaperone systems such as DnaKJE and GroEL, respectively (64). Thus, a rapid induction of the heat shock response upon detection of protein unfolding conditions is an essential element of the front-line defense of bacteria (65). Miwa *et al.* also provided evidence for an autoregulatory activity of IbpA, which occurs on post-transcriptional level (66). In the present study, we report AGXX®-induced IbpA-sfGFP foci formation in an AGXX® concentration-dependent manner, which is indicative of substantial protein aggregation in the cytoplasm of *E. coli*. We often observed the cellular localization of the IbpA-sfGFP foci at the cell poles, which aligns with observations of previous studies, where IbpA bound to denatured proteins is sequestered at the poles to relieve translation repression (27, 66, 67). Both our flow cytometry **(Fig. 3A)** and RT-qPCR analyses (**Supplementary Fig. S2**) point towards significant IbpA expression levels, which were only detectable 30 min after the beginning of the AGXX® treatment, which indicates that AGXX® stress occurs on a slower time scale compared to more fast-acting oxidants like RCS (38). Our data also suggest that AGXX® treatment impairs the integrity of the inner membrane, which could lead to a more uncontrolled uptake of silver ions or as previously shown, of drugs such as aminoglycoside antibiotics (20). DNA is one of the most important and therefore highly protected cellular biomolecules.

Microorganisms respond rapidly to DNA damage to protect themselves from mutations and/or to repair already damaged DNA. The upregulation of genes encoding SulA, a cell division inhibitor, and MutM, a DNA glycosylase, provided a first evidence of an activated DNA damage response in AGXX®-stressed *E. coli*. While AGXX® had been associated with proteotoxicity before, our present study is the first to show direct genotoxic effects of this antimicrobial: using the Gam-sfGFP biosensor, we provide evidence for a concentration-dependent increase in DNA double-strand breaks in living *E. coli* cells exposed to AGXX® (**Fig. 4**). Shee *et al.* have previously demonstrated that the bacteriophage Mu protein Gam detects DNA double-strand breaks with high specificity (28), which is due to the protein’s irreversible binding to DNA double-strand breaks. Moreover, DNA double-strand breaks, when bound by Gam, are no longer accessible to recombinases, proteins that are essential for the DNA damage repair (68–70). Surprisingly, pretreatment with the ROS scavenger thiourea did not significantly reduce Gam-sfGFP foci formation (*data not shown*), which suggests that the DNA damaging effect of AGXX® is likely not caused by ROS but instead could be a result of the direct interaction between DNA nucleobases and cationic Ag^+^ released from AGXX®. Interestingly, protein aggregation caused by AGXX® appears to occur faster than DNA damage.

A prominent signature of our RNAseq analysis of AGXX®-stressed CFT073 was the thiol-specific oxidative stress response (**Fig 7**). For instance, among the most upregulated genes during AGXX® stress were several encoding antioxidant systems, such as glutaredoxin 1 (*grxA*), thioredoxin 2 (*trxC*), superoxide dismutase (*sodA*), and alkyl hydroperoxidase reductase (*ahpF*). Moreover, we observed a strong induction of Cu^+^ response genes, including *cusRS* and the *cusFCBA* efflux system, indicating significant interference with the cellular metal homeostasis. Transcriptomic studies in AGXX®-treated *E. faecalis* also revealed the upregulation of Cu^+^ chaperone genes, which are export systems typically expressed in response to elevated Cu^+^ and Ag^+^ levels (16, 71). Highly induced *cusC* transcript levels, as detected in our RNAseq analysis, may therefore be a response to the possible influx of AGXX® microparticles or of silver ions released from AGXX® into the bacterial cytosol. Due to the similar coordination chemistry of Cu^+^ and Ag^+^ (72), it has been proposed that cells utilize redundant efflux systems to control the intracellular concentrations of both metals, which could explain why the Cu^+^ efflux systems were among the most strongly induced genes in our RNAseq analysis **(Fig 7A; Supplementary Table S4)**. Increased intracellular Ag^+^ concentrations as the main cause of DNA damage by AGXX® may also explain why the ROS scavenger thiourea did not alleviate DNA double-strand breaks, as Feng *et al.* showed that bacterial DNA is irreversibly damaged by Ag^+^ given that DNA replication was severely impaired even when Ag^+^-treated cells were recovered in fresh media in the absence of stress (11).

### ExPEC strains are better protected against the proteotoxic effects of AGXX®

Compared to the non-pathogenic *E. coli* K-12 strain MG1655, AGXX® treatments were less effective against ExPEC strain CFT073; we observed these differences on both phenotypic and macromolecular level (**Fig. 5&6**). Members of the UPEC pathotype may therefore employ additional and/or more sophisticated defense strategies to counter the antimicrobial effects of AGXX®. We have made very similar observations when UPEC strains were exposed to RCS, including treatments with hypochlorous acid (HOCl), the active ingredient of household bleach (30). *E. coli* have evolved numerous general and pathotype-specific mechanisms on both transcriptional and posttranslational level to fend off the toxic effects of antimicrobials such as RCS. General stress defense systems are encoded by genes located on the pan genome of *E. coli*. Consequently, these defenses are widely distributed among *E. coli* pathotypes, although their effectiveness may differ among *E. coli* pathotypes as we have reported for *rclC* (35). One of the most effective general defense systems is the conversion of ATP into polyphosphate, a chemical chaperone that protects *E. coli* from stressor-induced protein aggregation (46, 73, 74) (**Fig. 8**). Furthermore, molecular chaperones such as Hsp33, RidA, and CnoX are activated through thiol oxidation or N-chlorination, respectively (38, 62, 75–77). Another level of protection is provided through the transcriptional activation of general and pathotype-specific stress defense genes, which are directly controlled by stress-specific (in-)activation of transcriptional regulators (78). We previously identified one such UPEC-specific gene cluster as UPEC’s main defense system during severe RCS stress, enabling pathogen growth and survival at elevated RCS concentrations (30, 35). Expression of *rcrARB* is controlled by the redox-sensitive transcriptional repressor RcrR, which is inactivated by oxidation. While the precise biological function of *rcrA* and *rcrB* are still unknown, RcrB was shown to be exclusively responsible for UPEC’s superior resistance to RCS stress *in vitro* and phagocytosis, and *rcrB*-deletion strains were as sensitive to RCS as non-pathogenic *E. coli* strains that naturally lack this gene cluster. Interestingly, neither did we detect elevated *rcrB* transcript levels in response to AGXX®-stress nor was the *rcrB*-deficient strain more susceptible, suggesting different molecular defense mechanisms against AGXX® and RCS in UPEC. This is consistent with previous findings showing that the RcrR regulon was not expressed during exposure to H_2_O_2_, the main ROS generated by AGXX®, and that K-12 strain MG1655 and UPEC strains CFT073 and *rcrB* strains were equally resistant to this oxidant (30). In contrast to RCS, H_2_O_2_ is thiol-specific, orders of magnitude less bactericidal and therefore only kills bacteria after long exposure or at higher concentrations. Probably because bacteria generate this oxidant as an endogenous metabolic byproduct (79, 80), bacteria have evolved several efficient antioxidant systems to eliminate H_2_O_2_, including the peroxiredoxins AhpC and AhpF, whose expression was also highly induced in our RNAseq of AGXX®-treated CFT073 (**Fig. 7; Table S4**). Therefore, our data also suggest that AGXX® likely elicits a second stress mechanism other than ROS production, which UPEC appears to be better adapted to in comparison to K-12 strain MG1655. While these additional UPEC defense mechanisms are not identified quite yet, our data provide first insights into the cellular consequences induced by the unknown stressor: a comparison of changes in AGXX®-induced transcript levels between both strains revealed a reduced demand for molecular chaperones in UPEC, likely because this pathotype experiences less protein aggregation at AGXX® concentrations that kill MG1655. In our RNAseq analysis, we have identified 27 UPEC-specific uncharacterized genes among the significantly differentially expressed genes (**Supplementary Table S4&5**), 18 of which were highly upregulated. Further studies are now directed to investigate their potential contribution towards UPEC’s robust AGXX® defense.

### Bacteria produce polyP to protect themselves from AGXX®-induced damage

Gram-negative bacteria are known to produce long chains of polyP as a virulence strategy and to counter host defense mechanisms (42, 81–84). UPEC strains with defects in polyP production are characterized by their increased sensitivity to RCS (42) and elevated temperatures (43). This is attributed to polyP’s chaperone function, which protects the bacterial proteome from aggregation (43, 46, 47). Further, invasion of uroepithelial cells by UPEC is reduced in polyP-deficient bacteria (85) and mice infected with CFT073ΔpolyP display a lower bacterial load in the bladder (85), highlighting the important role of polyP in UPEC pathogenesis. Our growth and survival studies revealed a similarly protective effect of polyP against AGXX® (**Fig. 8).** Whether this is in response to AGXX®-generated ROS or potentially to silver ions that were released from the AGXX® formulation remains unclear given that ΔpolyP cells also showed increased susceptibility towards silver nitrate and hydrogen peroxide, respectively (**Supplementary Fig. S3**). While protection by polyP appears to be independent of the bacterial growth phase, the most significant reduction in survival was observed in stationary phase ΔpolyP cells **(Fig. 8)**. Our observation is consistent with previous studies, which revealed that bacteria lacking polyP are exquisitely sensitive to stress in stationary phase (86). The elevated susceptibility of polyP-deficient stationary phase cells is likely indirect and a result of the reduced expression of RpoS, the alternative sigma factor that is positively regulated by polyP (87–90). RpoS is responsible for the regulation of many genes that are required for stationary phase adaption, including the catalase-encoding gene *katE* as well as various stress defense genes that encode multidrug efflux pumps and antioxidant enzymes, of which some were upregulated in our RNAseq (91). Rao *et al.* have also demonstrated that the enzymatic activity of Ppk peaks in early stationary (92), providing additional support for the continuous accumulation of polyP during stationary phase growth. Exogenous addition of polyP to ΔpolyP cultures resulted in an almost complete rescue of the growth deficit, which could be due to increased RpoS expression. However, in light of our RNAseq data indicating significant imbalances in the metal homeostasis as well as based on the anionic nature of polyP and its established role as a metal chelator (93–95), it is also possible that exogenously added polyP protects polyP-deficient cells by chelating potentially released Ag^+^ ions **(Fig. 7**). PolyP has also been shown to reduce the mutation rate of bacterial DNA and protect from DNA damage-induced cell death (96), which could explain the increased susceptibility of the ΔpolyP strain. Likewise, it has been reported that polyP acts as a metal chelator and inhibitor of the Fenton reaction (97), which may help cells to protect from AGXX® stress. Given the various roles of bacterial polyP, it is not surprising that many pathogens rely on protection by polyP, making the bacteria-specific enzyme Ppk1 an ideal drug target. Several efficient inhibitors of Ppk1 have been identified, including mesalamine, an FDA-approved drug used to treat ulcerative colitis (98), and gallein (73, 99, 100). Treatment with either of these inhibitors severely compromises bacterial survival during oxidative stress, biofilm-formation, and colonization. Independent studies have confirmed the Ppk1 inhibitory effects of mesalamine and gallein (46, 101–103) and provided evidence that oxidative stress defense systems like polyP positively affect pathogen colonization in the host and negatively affect the innate immune response (85, 100, 104, 105). Thus, targeting processes such as polyP production, which are only essential for bacterial survival in the context of infections and directly contribute to bacterial virulence have the potential to further sensitize UPEC towards antimicrobial agents such as AGXX® (4, 106), which we aim to test in the future.

## MATERIALS AND METHODS

### Bacterial strains and growth conditions

All strains and oligonucleotides used in this study are listed in the **Supplementary Table S6**. Unless otherwise stated, overnight bacteria cultures were grown aerobically in Luria Bertani (LB) broth (Millipore Sigma) at 37 °C and 300 rpm. For subsequent assays, overnight cultures were diluted into 3-(N-morpholino) propanesulfonic acid minimal media containing 0.2% glucose, 1.32 mM K_2_HPO_4_, and 10 µM thiamine (MOPSg) (107) and incubated at 37 °C under shaking conditions.

### Preparation of AGXX® formulations

Largentec GmbH (Berlin, Germany) developed and provided all AGXX® microparticles used in this study. Briefly, the AGXX® formulations 394C and 823 are composed of silver powders ranging in particle size from 1.5-2.5 µm (MaTeck, Germany). In contrast, silver powders with particle sizes greater than 3.2 µm (Toyo, Japan) were utilized for 383. Silver powder coating was meticulously applied to hollow glass microparticles, followed by coating with Ru (III) ions. Subsequently, Ru (III) ions underwent oxidation to RuO_4_ via sodium hypochlorite. The addition of sodium nitrite reduced RuO_4_ to Ru. Afterward, the AGXX surface was conditioned with 50 mM ascorbate for an extended time. It was then filtrated, rinsed with deionized water, and dried with a hot air blower.

### Growth curve-based assays

Overnight cultures of the indicated strains were diluted ∼25-fold into fresh MOPSg and grown at 37°C under shaking conditions until the mid-log phase (OD_600_= ∼0.5) and cultivated either in the presence or absence of increasing concentrations of the indicated AGXX® formulations. Absorbance (*A*_600nm_) was recorded every 30 min for 4 hrs using the spectrophotometer (Biomate 3, Thermo Scientific). For growth-curve-based studies in the Tecan Infinite plate reader, overnight cultures of the indicated strains were diluted ∼25-fold into fresh MOPSg, grown at 37 °C until late logarithmic phase (OD_600_= ∼2), diluted to an OD_600_= 0.05 and cultivated in the presence of the indicated AGXX® concentrations. For supplementation with exogenous polyP, 4 mM polyP was added to the Δ*ppk* cultures.

### Bacterial survival assays after AGXX® exposure

Overnight cultures were diluted into fresh MOPSg media to an OD_600_=∼0.05 and cultivated until OD_600_ of 0.3 was reached before being transferred into 125ml sterile flasks and grown in the presence or absence of increasing concentrations of s AGXX®. For the ROS quenching experiments, cultures were pretreated with 70mM thiourea 60 minutes prior to AGXX®. At the indicated time intervals, cells were serially diluted in PBS (pH 7.4) and spotted on LB agar for CFU counts after overnight incubation. Survival percentages were calculated as the ratio of CFU_treated_ / CFU_untreated_ samples.

### Intracellular ROS measurements

The redox-sensitive, cell-permeant dye 2′,7′-dichlorodihydrofluorescein diacetate (H_2_DCFDA) (Thermo Fisher Scientific) was used to quantify intracellular ROS levels. Mid-log CFT073 cultures were either left untreated or treated with the indicated concentrations of AGXX®394C for 60mins. Samples were normalized to an OD_600_ = ∼ 1.0, washed twice in PBS, resuspended in prewarmed PBS containing 10 µM H_2_DCFDA, and incubated in the dark at 37°C. After 30min, samples were washed twice in PBS and DCF fluorescence measured at excitation/emission (exc./em.) wavelengths of 485/535nm in a Tecan 200 plate reader. Cells were pretreated with 70 mM thiourea for cellular ROS quenching before the addition of AGXX®. 5 mM Paraquat was included as a positive control.

### Quantification of hydrogen peroxide (H_2_O_2_)

The fluorescent probe Amplex^TM^ Red (Invitrogen) was used to quantify the generated cellular H_2_O_2_ levels. Exponentially growing cells were either left untreated or treated with AGXX®394C for 60 minutes. Cells were washed twice in PBS, stained with the Amplex-red HRP working solution as instructed by the manufacturer, and incubated in the dark at 37°C for 30mins. Resorufin fluorescence was measured at exc./em. wavelengths of 550/590nm in a Tecan 200 plate reader. 12mM H_2_O_2_ was included as a positive control.

### Quantification of superoxide

Dihydroethidium (DHE) (Invitrogen) was used to monitor intracellular levels of superoxide. Exponentially growing cells were either left untreated or treated with AGXX®394C for 60 min. Cells were harvested, washed, and resuspended in PBS (pH 7.4) to an OD_600_ = 1.0, followed by incubation at 37 °C with 50µM DHE for 60 min under shaking conditions before fluorescence was measured at exc./em. wavelengths of 518/606nm in a Tecan 200 plate reader. 5 mM Methyl viologen was included as a positive control.

### Gene expression levels using qRT-PCR

Overnight cultures of the indicated strains were diluted into MOPSg media to an OD_600_=0.08 and cultivated until the mid-log phase (OD_600_=0.3). Cells were either left untreated or treated with AGXX®. At the indicated time points, transcription was stopped by the addition of ice-cold methanol, total RNA extracted using a commercially available RNA extraction kit (Macherey-Nagel), remaining genomic DNA removed using the TURBO DNA-free kit (Thermo Scientific), and cDNA synthesized using the PrimeScript cDNA synthesis kit (Takara). qRT-PCR reactions were set up according to the manufacturer’s instructions (Alkali Scientific). Transcript levels of the target genes were normalized to the 16S rRNA-encoding *rrsD* gene, and relative fold changes in gene expression were calculated using the 2^-ΔΔCT^ method.

### Membrane permeability assessment

PI uptake was used to determine plasma membrane integrity following AGXX® exposure. Exponentially growing CFT073 cells were either left untreated or treated with the indicated AGXX®394C concentrations for 60 min. Samples were harvested, washed twice, and resuspended in PBS (pH 7.4) at an OD_600_=0.5. PI (Thermo Fisher Scientific) was added to a final concentration of 0.5 µM, and samples were incubated in the dark for 30 min. Fluorescence was measured at exc./em. wavelengths of 535/617nm. Samples exposed to 4 µg/ml polymyxin B were included as a positive control.

### Live/Dead Staining

The experiment was performed as previously described (20, 108). Briefly, mid-log cultures of CFT073 were either left untreated or treated with 40 µg/ml AGXX394C for 60 min. Cells were harvested, washed twice, and resuspended in PBS (pH 7.4) at an OD_600_ =0.2. Samples were incubated with 6 µM dyes SYTO9 and 30 µM PI for 15 mins in the dark at room temperature. Cells were transferred onto a glass slide and covered with a 1% agarose pad prior to visualization using a Leica SP8 Confocal system equipped with a DMi8 CS inverted microscope. Polymyxin B-treated cells were included as positive controls.

### Expression and microscopy of GamGFP foci

Overnight cultures of MG1655-Gam-sfGFP were cultivated in LB in the presence of 200 ng/ml doxycycline to induce Gam-sfGFP. Cultures were then diluted into doxycycline-containing MOPSg to an OD_600_= ∼0.08 and grown until mid-log phase (OD_600_= ∼0.3). Cells were then left untreated or treated with the indicated concentration of AGXX® or ciprofloxacin for 3h. Cells were then washed twice, resuspended in fresh PBS, incubated with 10 µg/ml DAPI and 5 µg/ml FM4-64 in the dark for 15 minutes, washed again twice, and resuspended in PBS. Cells were imaged on 1% agarose gel pads via fluorescence microscopy using a Leica SP8 confocal system equipped with a DMi8 CS inverted microscope. 25 µg/ml Ciprofloxacin, a known DNA-damaging antibiotic, was included as a positive control.

### Extraction and visualization of Protein Aggregate after AGXX stress

The experiments were described following the procedure described before (37).

### IbpA-sfGFP expression and binding to protein aggregates *in vivo*

Overnight cultures of the indicated strains were diluted into fresh MOPsg to an OD_600_=∼ 0.05 and grown to mid-log phase (OD_600_= ∼0.3), followed by AGXX® treatment for indicated time points. For flow cytometry analysis of IbpA-sfGFP expression, sample volumes were normalized to an OD_600_=0.05 in PBS (pH 7.4) prior to analysis in the flow cytometer (BD FACS Melody) using the FITC channel. At least 10,000 events were recorded, and figures were generated using FCSalyzer. For visualization of IbpA-sfGFP binding of protein aggregates in the cell, samples were washed twice, resuspended in fresh PBS and subsequently stained with 5 µg/ml DAPI and 5 µg/ml FM4-64 in the dark for 15 min. Cells were washed twice, resuspended in PBS, and then imaged on 1% agarose gel pads and via fluorescence microscopy using a Leica SP8 confocal system equipped with a DMi8 CS inverted microscope. At least 100 cells per independent experiments were counted blindly.

### RNAseq analysis, Differential Gene Expression and Data visualization

Samples of AGXX®-treated and untreated CFT073 cells were collected as described for qRT-PCR. After extraction of total RNA (Macherey & Nagel) and removal of the residual DNA using the TURBO DNA-free kit (Thermo Scientific), rRNA was depleted using the Illumina Ribo Zero Kit (Illumina) for Gram-negative bacteria. A total of 150 bp single-end sequencing was performed on an Illumina HiSeq 2500 by Novogene (Sacramento, USA). Differential gene expression analysis of three biological replicates, including normalization, was performed in the bioinformatics platform Galaxy (109). Briefly, RNAseq reads were mapped to the CFT073 reference sequence ([GCA_000007445.1](https://www.ncbi.nlm.nih.gov/bioproject/313)) using HISAT2 (110). Then, the number of reads mapped to each gene was counted using featureCounts (111). Finally, differential gene expression was visualized using DESeq2(112) with an adjusted *P* value cut off *P* ≤ 0.05 and log_2_FC cut off = 1.5. The evaluation of the differential gene expression was further visualized as a M/A plot where *M-*values (log_2_ fold change values) were plotted against A-values (log_2_ base mean). Using the KEGG database, gene IDs were used to search for biological processes and grouped under broader Gene Ontology (GO) terms. Statistically significant DEGs found to have more than one biological process were sorted and categorized accordingly. Finally, GO terms were plotted against the total number of DEGs in a bar graph.

## Statistical analyses

All statistical analyses were performed in GraphPad Prism version 8.0.

## ACKNOWLEDGEMENT

This work was supported by the NIAID grants R15AI164585 and 1R03AI174033-01A1 and the Illinois State University Faculty Research Award (to J.-U. D.). We thank members of the Dahl lab and the lab of Dr. Kyle Floyd (Illinois State University) for feedback and proofreading the manuscript. The Largentec GmbH team is acknowledged for providing the different AGXX® formulations and for helpful discussions.

## SUPPLEMENTARY INFORMATION

**Supplementary Fig. S1:**
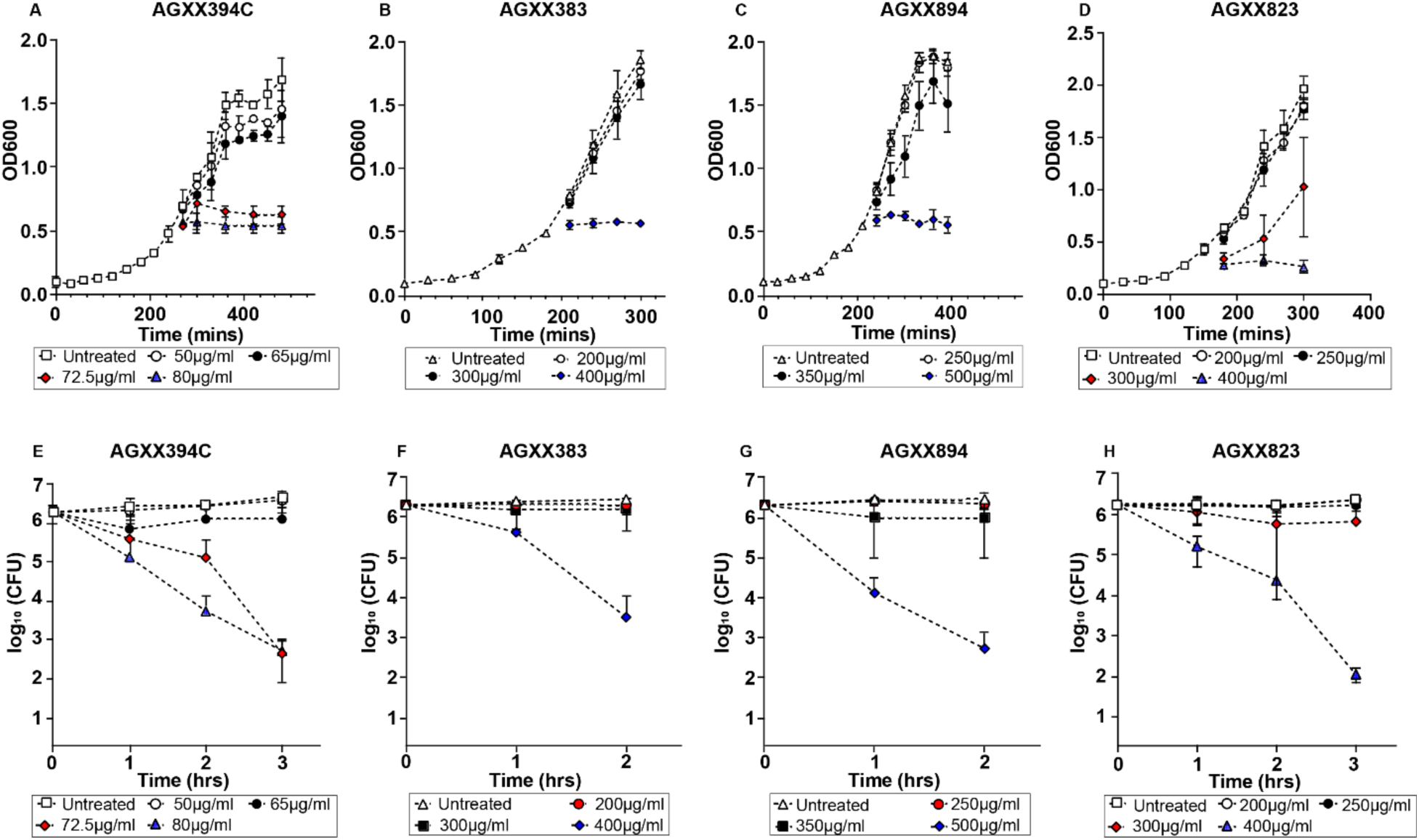
AGXX® formulations differ in their antimicrobial activities. Growth and survival studies were performed in UPEC strain CFT073, which was grown in MOPSg media to mid-log phase and treated with the indicated concentrations of AGXX®394C (**A; E**), AGXX®383 (**B; F**), AGXX®894 (**C; G**), and AGXX®823 (**D; H**), respectively. (**A-D**) Absorbance at 600 nm (OD_600_) was recorded every 30 mins for 4 hrs (n=3-4, ±S.D.). (**E-H**) For assessment of bacterial killing, samples were taken every 60 minutes and serially diluted in PBS. Five µl of serial dilutions were spotted onto LB agar for colony forming units (CFU) counts after overnight incubation (n=4-7, ±SD)

**Supplementary Fig. S2:**
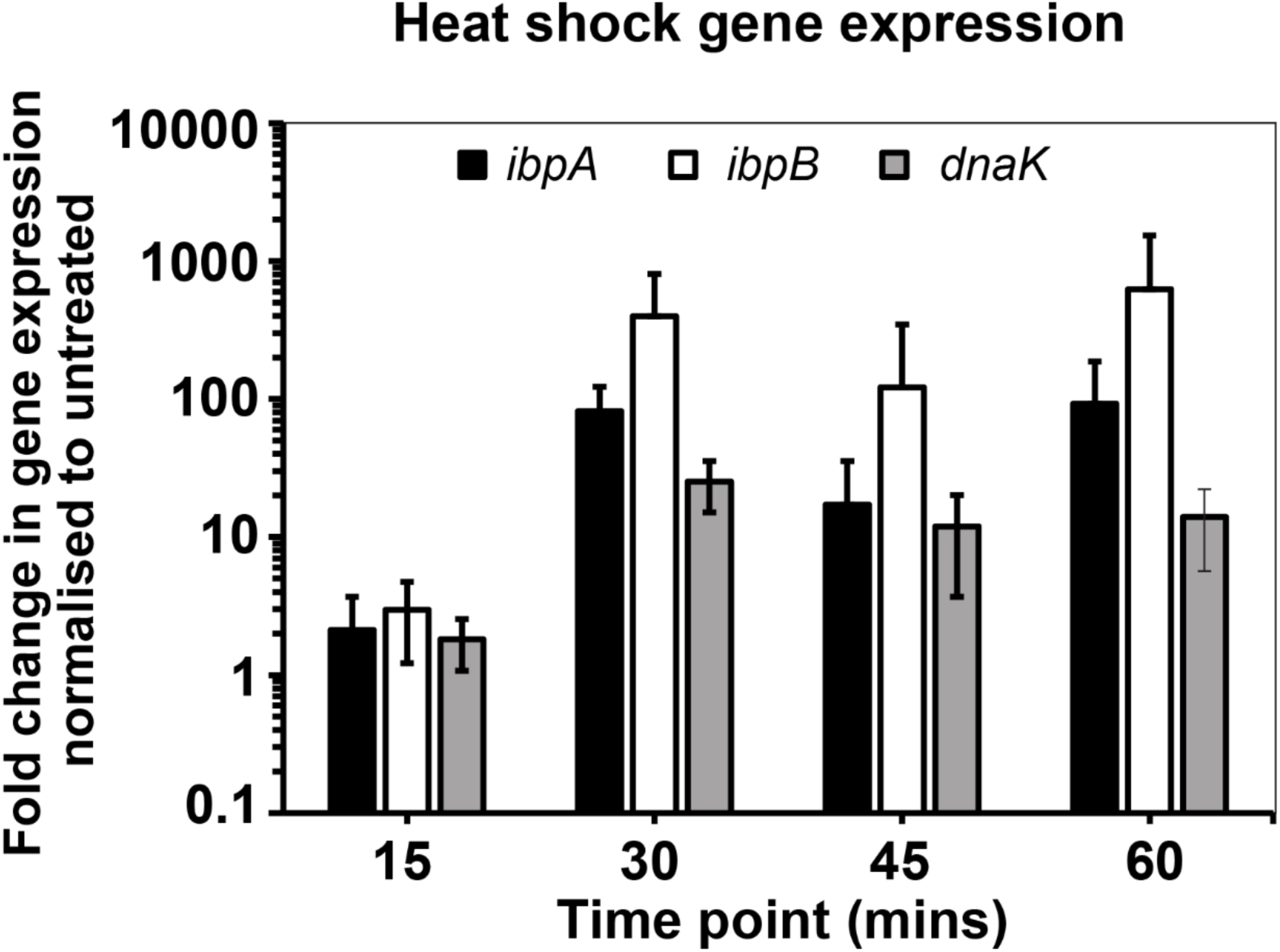
Time-course of the transcriptional response of UPEC to AGXX® treatment. Exponentially growing CFT073 cultures were exposed to 250 µg/ml AGXX®823 and samples collected at the indicated timepoints for RNA extraction, removal of genomic DNA, and reverse transcription of mRNA into cDNA. qRT-PCR analysis was performed for select heat shock genes and normalized to the housekeeping gene *rrsD* and untreated samples (n= 4, ±S.D.)

**Supplementary Fig. S3:**
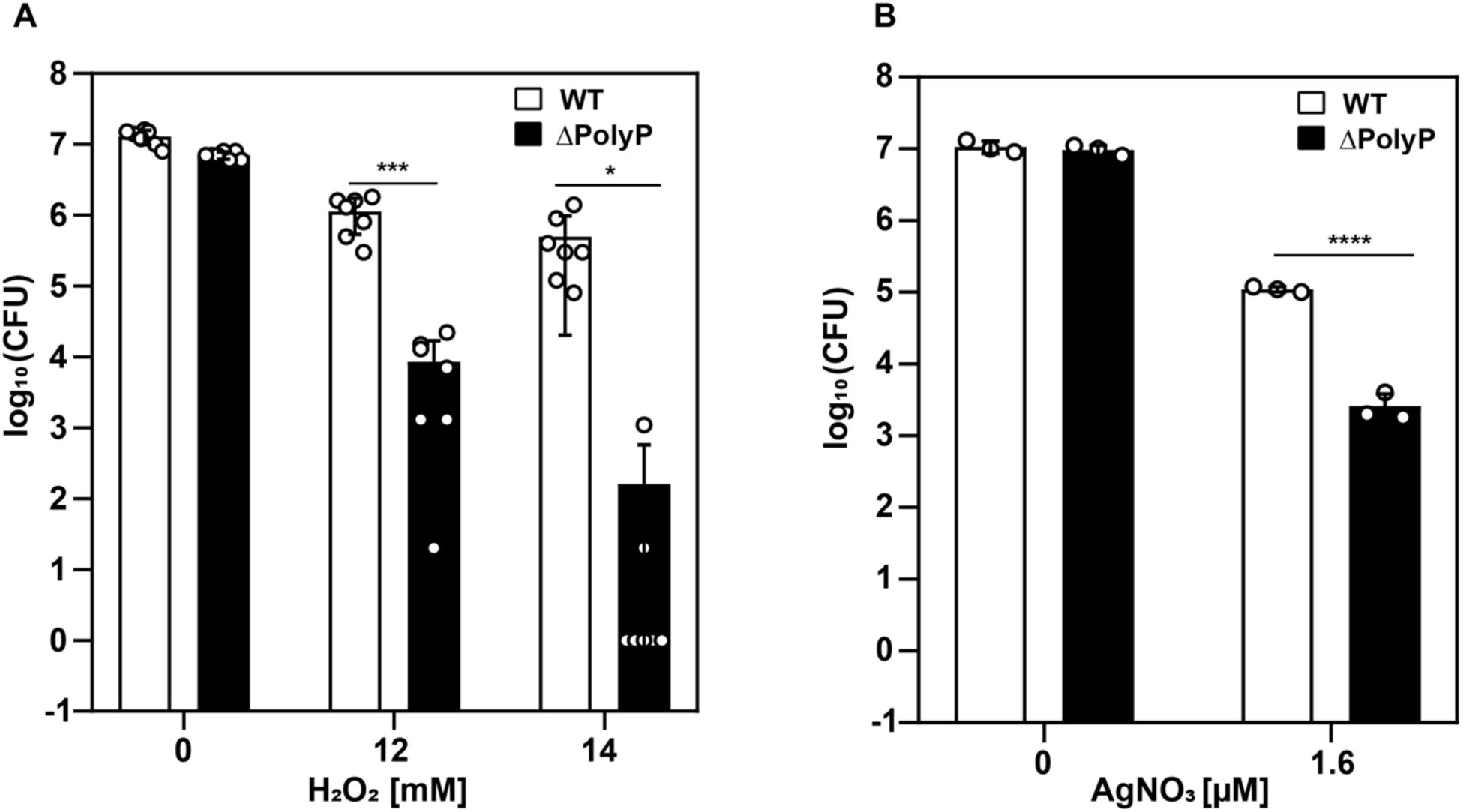
Polyphosphate protects UPEC from hydrogen peroxide and silver. Exponentially growing CFT073 and ΔpolyP cells were exposed to the indicated concentrations of hydrogen peroxide (**A**) and silver nitrate (**B**) for 180 min before samples were serially diluted in PBS, spot-titered on LB agar, and incubated for 20 hrs for CFU counts (n= 3-6, ±S.D.). student t-test; ns = *P* > 0.05, * *P* < 0.05, *** *P* < 0.001, **** *P* < 0.0001.)

